# Gla-domain mediated targeting of externalized phosphatidylserine for intracellular delivery

**DOI:** 10.1101/2022.06.13.495901

**Authors:** Jonathan Hardy, Maxine Bauzon, Francis Blankenberg, Masamitsu Kanada, Ashley Makela, Charles K.F. Chan, Christopher H. Contag, Terry Hermiston

**Affiliations:** Department of Pediatrics, Stanford University School of Medicine, Stanford, California, United States of America; Biologics Research US, Bayer HealthCare, San Francisco, California, United States of America; Department of Radiology/MIPS, Stanford University, Stanford, California, United States of America; Department of Surgery, Stanford University, Stanford, California, United States of America; Department of Microbiology and Immunology, Stanford University School of Medicine, Stanford, California, United States of America

**Keywords:** phosphatidylserine, extracellular vesicles, GLA-domain, stem cells, cancer, infection

## Abstract

Phosphatidylserine (PS) is a negatively charged phospholipid normally localized to the inner leaflet of the plasma membrane of cells but is externalized onto the cell surface during apoptosis as well as in malignant and infected cells. Consequently, PS may comprise an important molecular target in diagnostics, imaging and targeted delivery of therapeutic agents. While an array of PS binding-molecules exist, their utility has been limited by their inability to recognize PS and internalize diagnostic or therapeutic payloads. We describe the generation, isolation, characterization, and utility of a PS binding motif comprised of a carboxylated glutamic acid (GLA) residue domain, that both recognizes cell surface-exposed PS and is internalized into these cells after binding. Internalization is independent of the traditional endosomal-lysosomal pathway, directly entering the cytosol of the target cell in a rapid and energy-independent fashion. We demonstrate that this PS recognition extends to extracellular vesicles and stem cells and that GLA-domain conjugated probes can be detected upon intravenous administration in animal models of infectious disease and cancer. GLA domain binding and internalization offers new opportunities for targeting specific cells for imaging and delivery of therapeutics.

## Introduction

Phosphatidylserine (PS) is a negatively charged phospholipid normally localized to the inner leaflet of the plasma membrane of cells. Exposure of PS on the surface of apoptotic cells is a normal physiological process that triggers their rapid removal by phagocytic engulfment under noninflammatory conditions via receptors primarily expressed on immune cells (1). Importantly, PS is also externalized on many diseased cells (e.g. cancer and infections with viral, bacterial, and parasitic agents) where these cells take advantage of the evolutionarily conserved global immunosuppressive signal to evade immune detection and clearance (2). Due to the large number of cellular processes conferred by cell-surface expression of PS, there has been significant interest in developing PS-binding molecules or PS-recognizing antibodies as imaging probes and therapeutic agents for disease detection and treatment (3, 4).

While Annexin V is the best studied of the PS binding proteins due to its widespread use in cell biology and clinical experience (5), the ability to manipulate this protein for therapeutic utility has been significantly challenged by the need for trimerization to interact with PS. An alternative agent for PS targeting are the GLA (gamma carboxyglutamic acid) domains of proteins. GLA domains are relatively short (~45 amino acid) polypeptide segments that are rich in glutamate residues carboxylated by a vitamin K-dependent post-translational modification (6, 7). GLA domains appear in approximately two dozen complex human proteins, many of which play a critical role in blood coagulation. The coagulation factors Prothrombin, Factors VII, IX, and X, and anti-coagulant Proteins C, S, and Z all contain GLA domains that allow them to assemble in a Ca^++^-dependent fashion onto the PS-rich surface of activated platelets and affect blood coagulation. Because these are naturally occurring and relatively small (<100 amino acid residues), compared to other PS-binding entities, such as annexin or antibodies, and may lack the need for dimer or trimer formation, we have begun isolating the GLA domains from the multi-functional parent proteins and engineering these as a PS-targeting platform for imaging and drug delivery. Using annexin V as a well-described industry standard PS binding protein for comparison, we demonstrate that the GLA domain can be isolated in a fashion that is capable of distinguishing PS expressing cells *in vitro* and *in vivo*, and moreover that these GLA domain proteins, and attached cargo, are internalized into cells.

The structure of GLA-domain containing proteins (Fig 1A) reveals their modular nature, with a prototypical n-terminus GLA domain, EGF (or Kringle-like) elements, and a catalytic/receptor binding domain at the carboxy terminus of the protein. Fig 1B shows a comparison of the amino acid sequences from GLA domains of seven coagulation system proteins, highlighting the conserved cysteine residues and characteristic carboxylated glutamic acid residues (GLA). From among these proteins, we have chosen the GLA domain of protein S to test the development of GLA domains as a platform for diagnostic and therapeutic applications. Native human protein S (8, 9) was first identified as a glycoprotein which played a role in anticoagulation (10). Since then, its importance in a number of other functions has been identified. Protein S contains an amino-terminal GLA domain, four EGF-like domains, and dual carboxy-terminal domains that combine to bind immunosuppressive Tyro-3/Axl/Mer receptors (TAMRs) expressed on phagocytes. In order to study the binding properties of the GLA domain of protein S, we deleted the TAMR-binding domains, leaving the EGF-like domains and the GLA domain intact. In addition, a 6xHis-tag was added to facilitate purification and detection (Fig 1C). This construct was designated “GlaS”, to specify its origin. Here we demonstrate GlaS binding and internalization into cells with PS on their surface as an indicator of malignancy, infection, and stress.

**Figure 1.**
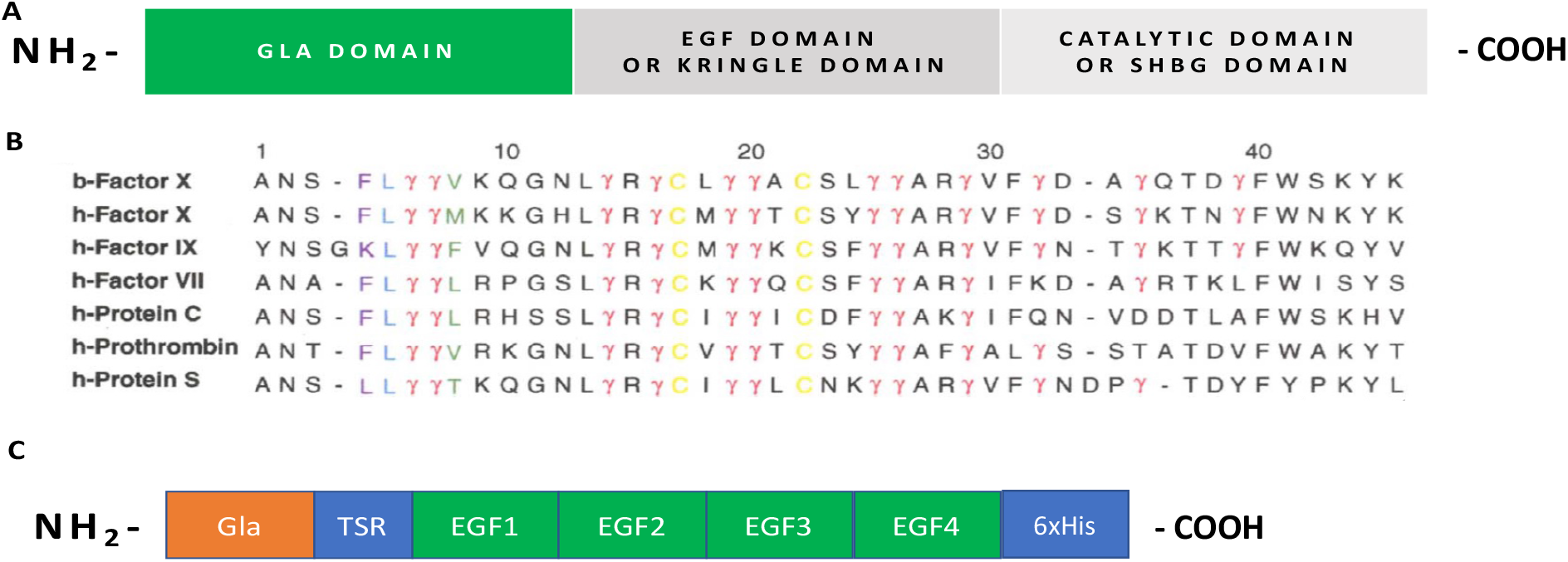
Coagulation factors, their associated GLA domains and the engineered prototype of ProS. A) Diagram of a prototypical coagulation factor with an N-terminal GLA domain followed by an EGF or Kringle domain region, and a catalytic or SHBG carboxy terminal end; B) GLA domain amino acid sequence comparison of coagulation factors, roughly 45 amino acids with conserved cysteine residues and the 9-12 gamma-carboxylated glutamic acid residues embedded within the GLA domain, indicated by γ, and giving these proteins their name; C) Protein S GLA and EGF domain construct (GlaS) with the N terminal GLA domain, thrombin sensitive region (TSR), four EGF domains. The carboxy terminal SHBG domain was deleted and a 6xHis-tag was added to facilitate purification and detection.

## Materials and methods

### GlaS construction and production

DNA encoding the first 283 amino acids of human Protein S with the addition of a 6x His tag was synthesized at GenScript (Piscataway, NJ). This insert contained the signal peptide, propeptide, GLA and 4 EGF domains of Protein S flanked by AgeI and NotI restriction sites and the expressed protein was called GlaS. The insert was sequence verified and subcloned into pEAKflCMV (Eagle Scientific vector modified to include full length CMV enhancer promoter sequence) using AgeI and NotI resulting in pEAKflCMV Protein S GLA+EGF.

### Animals

Eight-week-old female CD1 and BALB/c mice were purchased from Charles River Laboratories, and the mice were housed and maintained in standard BSL-2 conditions. The experiments conducted with bacterial pathogens were conducted according to animal protocol number 11601, and with tumors, protocol number 28411, both approved by the Administrative Panel for Laboratory Animal Care and the Institutional Animal Care and Use Committees at Stanford University. Mice were housed in ventilated cages in climate-controlled facility and provided with food and water ad libitum. Nestlets were provided. The mice were anesthetized with 2% isoflurane in oxygen during mammary fat pad injection of 4T1 tumor cells. Isoflurane was not employed for standard tail vein injection of *Listeria*. Mice infected with *Listeria* or injected with tumor cells were monitored daily for signs of distress such as matted fur, hunched posture, pale skin color and lethargy. None of these signs were observed in any animals during these studies nor did any of the tumors reach 1 cm in size which would have necessitated euthanasia as per IACUC guidelines. The animals were humanely euthanized with CO_2_ for dissection and sample collection. No animals died prior to euthanasia for sample collection.

### Protein Labeling and Cell Staining

For fluorescence, conjugation of Cy5 and FITC was achieved using Amersham (GE Heathcare) and Molecular Probes (Invitrogen) labeling kits, respectively, according to the manufacturer instructions. Human MDA-MB-231 and MCF7, murine 4T1 and MET-1, and COS-1 monkey kidney cells were used to test fluorescence staining of labeled GlaS and annexin V. The cells were plated in 24-well plates at 6×10^4^ cells per well or Eppendorf chamber slides at 1×10^4^ cells per well, and apoptosis was induced the next day, using 2mM H2O2, or 0.2mM tertiary-Butyl hydroperoxide (t-BHP) for time points from 30 min to 2 hrs. After induction, the wells were washed with Annexin Binding Buffer (AB; Santa Cruz Biotech) and stained with labeled protein. From past experience and literature, 5.5 ug/ml of annexin V protein was used for staining. This amount was adjusted for equimolar addition of GlaS by assuming the molecular weights of annexin to be 36 kD and the recombinant GlaS to be 30 kD. The cells were stained for 5-30 min. It was determined that less than 5 min was sufficient for GlaS staining (data not shown). Hoechst 33342 dye was used for visualizing nuclei. The wells were then washed with AB and observed using the EVOS fluorescence microscope (Life Technologies) while still viable. For confocal microscopy, a Leica TCS SP8 confocal microscope in the Stanford Cell Sciences Imaging Facility was employed. For toxicity studies, GlaS was added to trophoblast stem cells (TSCs) and cell viability tested with trypan blue using a Nexcelom Cellometer (Lawrence, MA).

### Extracellular Vesicle (EV) Isolation

EV-depleted medium was prepared by 18 h-ultracentrifugation of fetal bovine serum (FBS) at 100,000g, 4 °C (11). 4T1 cells were seeded at 1 × 10^6^ cells in 100 mm cell culture dishes and cultured in 10 mL of the EV-depleted media for 2 days, and EVs were harvested as described previously (12). Briefly, conditioned medium was centrifuged at 600g for 30 min to remove cells and debris. The supernatant was centrifuged again at 2,000g for 30 min to remove apoptotic bodies (13). EVs were collected by ultracentrifugation at 20,000g for 60 min using Optima XL-90 Ultracentrifuge and 90Ti rotor (Beckman Coulter).

### Cancer, Infection and Imaging of Mice

To test the labeled proteins for the ability to detect tumors, 5×10^4^ 4T1-luc cells were implanted into groups of 5 male BALB/c mice, in the left axillary fat pad. Tumor location and size were assessed using *in vivo* bioluminescence imaging (BLI) of mice daily, starting at 1 week post-implantation. The mice were then treated on day 11 post implantation with 13 mg/kg body weight of intraperitoneal (IP) doxorubicin, and 300 mg/kg body mass of Cy5-GlaS. BLI was performed the next day. Control tumor-bearing mice were left untreated.

To test for the specific localization of fluorescent GlaS to infection in live animals, CD1 mice were injected intravenously with bioluminescent *Listeria monocytogenes*. This bacterial pathogen infects many organs including the spleen, in which extensive apoptosis of monocytes and granulocytes occurs. Following infection, mice were imaged daily. When splenic signals were evident (day 2 post infection for 2 × 10^5^ colony forming units of bacteria in 8 week-old CD1 female mice), 300 mg/kg body mass of Cy5-GlaS was injected into mice, the animals were sacrificed 30 min later, and the spleens removed, frozen in OCT, and sectioned for fluorescence microscopy. Uninfected control mice were also assayed by the same methodology.

### Flow Cytometry and Microscopy of HSC

Freshly labeled FITC-GlaS, prepared as described above, was employed. Murine hematopoietic stem cells (HSCs) were purified from normal mouse bone marrow by staining for c-Kit+, sca+, lineage-negative cells. To further characterize the cells, SLAM marker staining was also performed (14). These markers stain long term HSCs that self-renew and differentiate to all lineages of the hematopoietic system. Subsequent staining with FITC-GlaS revealed over 90% percent positive in SLAM-staining cells, as shown in Supplemental Figure 3. The cells were then sorted for FITC and examined with confocal microscopy, using 10 μg/mL Hoechst 33342 (Life Technologies) for nuclear visualization (Supplemental Figure 3d).

## Results

A variety of diseased cells have been described to express PS on their surface (2). To validate that the GlaS protein, as engineered, retains PS recognition activity, we briefly treated human cells with a low concentration of peroxide to induce cell surface PS expression. Low peroxide levels and brief incubation times were used in order to observe still viable cells with minimal PS exposure, and both GlaS and annexin V were tested for binding to the PS-expressing cells. In this system we were able to demonstrate binding of FITC-labeled GlaS (Fig 2A), similar to FITC-annexin on peroxide treated MDA-MB-231 cells (Fig 2C). Binding was also observed in peroxide treated human MCF-7 (Fig 2D), murine MET-1 (Fig 2E), and murine 4T1 cells (Fig 2F). Within the parameters of the assay, no significant FITC-GlaS or FITC-annexin staining was observed on untreated cells (Fig 2B). When GlaS and annexin V were used in co-staining experiments, the two proteins did not always bind the same cells (Fig 3). Although there was overlap, some cells clearly stained only with annexin V (Fig 3A) or GlaS (Fig 3B). In addition, GlaS bound in the presence of 1,000-fold molar excess of annexin V (Fig 3C), suggesting that these two PS-binding proteins interact with PS in a distinctive manner.

**Figure 2.**
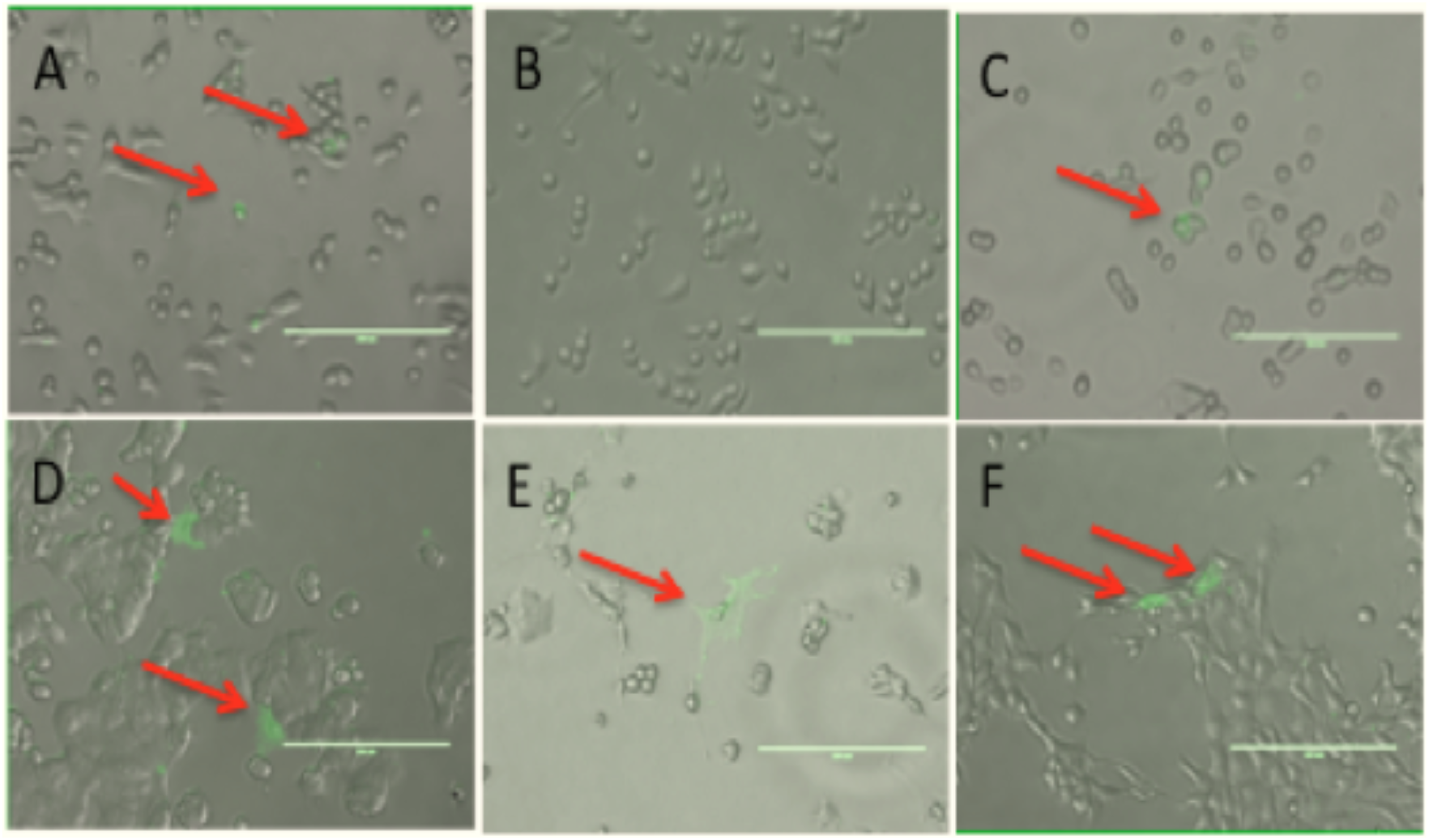
GlaS and annexin V staining of human and murine breast cancer cell lines treated with peroxide to induce PS externalization. A) Human MDA-MB-231 cells treated with peroxide and stained with FITC-GlaS.; B) Untreated MDA-MB-231 cells stained as in A; C) Treated MDA-MB-231 cells stained with annexin V; D) Human MCF-7 cells treated with peroxide and stained with GlaS; E) Murine MET-1 cells, as in D; F) Murine 4T1 cells, as in D.

**Figure 3.**
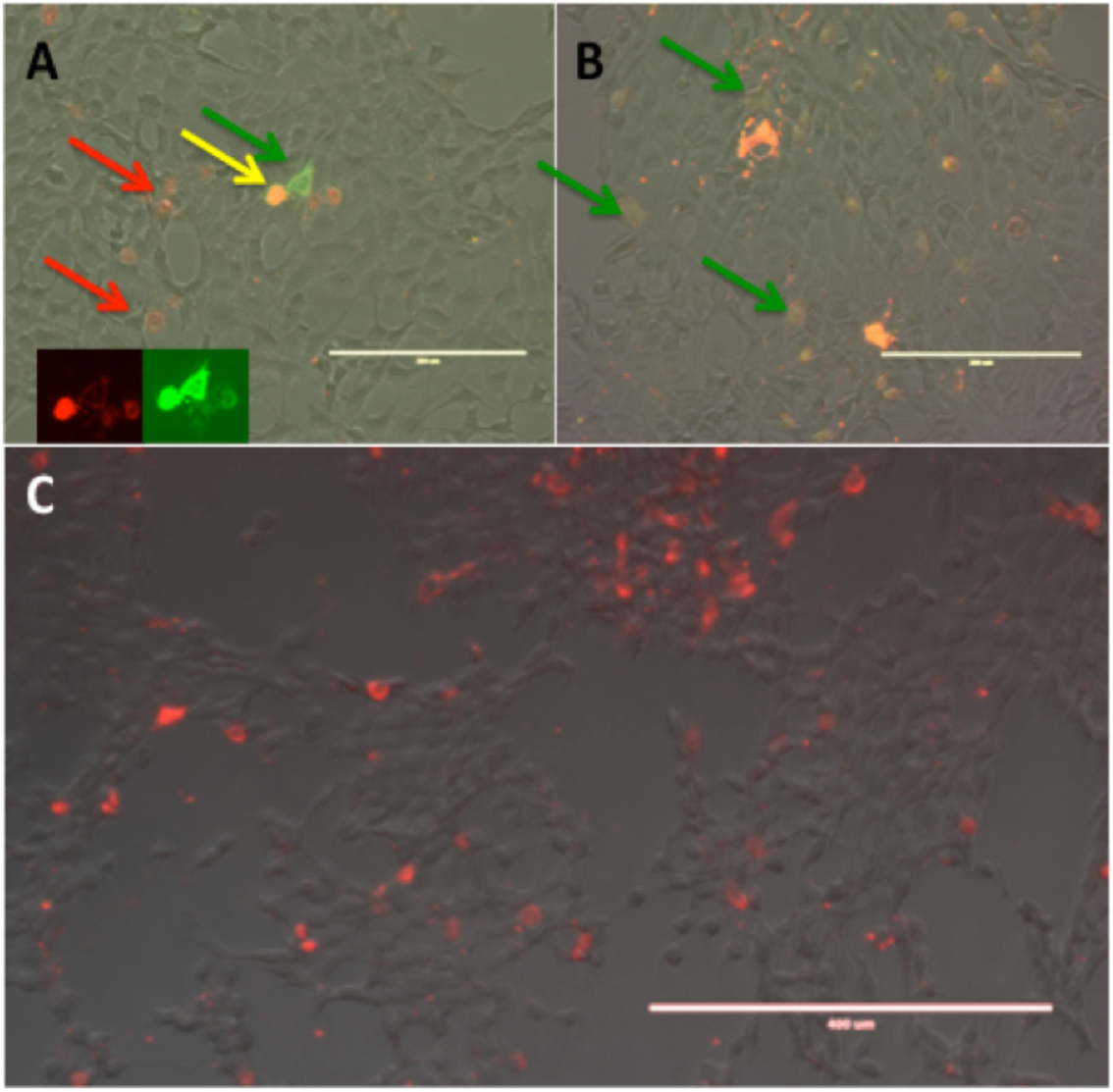
Overlapping, yet distinct, cellular localization of GlaS and annexin V. A) murine 4T1 cells treated with peroxide and stained with Cy5-GlaS (red) and FITC-annexin V (green). Yellow arrow, co-localized signals; red arrows, cells staining with GlaS and not annexin V; green arrow, cell staining relatively brighter with annexin V but less bright with GlaS, indicating distinct binding patterns (inset shows GlaS and annexin staining separately); B) treated 4T1 cells stained with FITC-GlaS (green) and Cy5-annexin V (red), green arrows indicate cells staining with GlaS and not annexin V; C) Cy5-GlaS (red) staining of treated 4T1 cells pre-incubated with 1,000-fold excess of unlabeled annexin V.

To further explore the differential interaction between FITC-GlaS and Cy5-annexin V on PS-expressing cells, 4T1 cells were treated briefly with peroxide, stained with labeled GlaS and annexin V, and the live cells were imaged with confocal microscopy (Fig 4). In many cells, GlaS was internalized, whereas annexin V was not (Fig 4A and B). The converse was not observed in any cells, *i*.*e*., internalized annexin V together with surface-localized GlaS. As an internal control, we observed vesicles, among these cells, that were of a size consistent with apoptotic bodies (1-5 μm) that stained with both GlaS and annexin V (Fig 4C).

**Figure 4.**
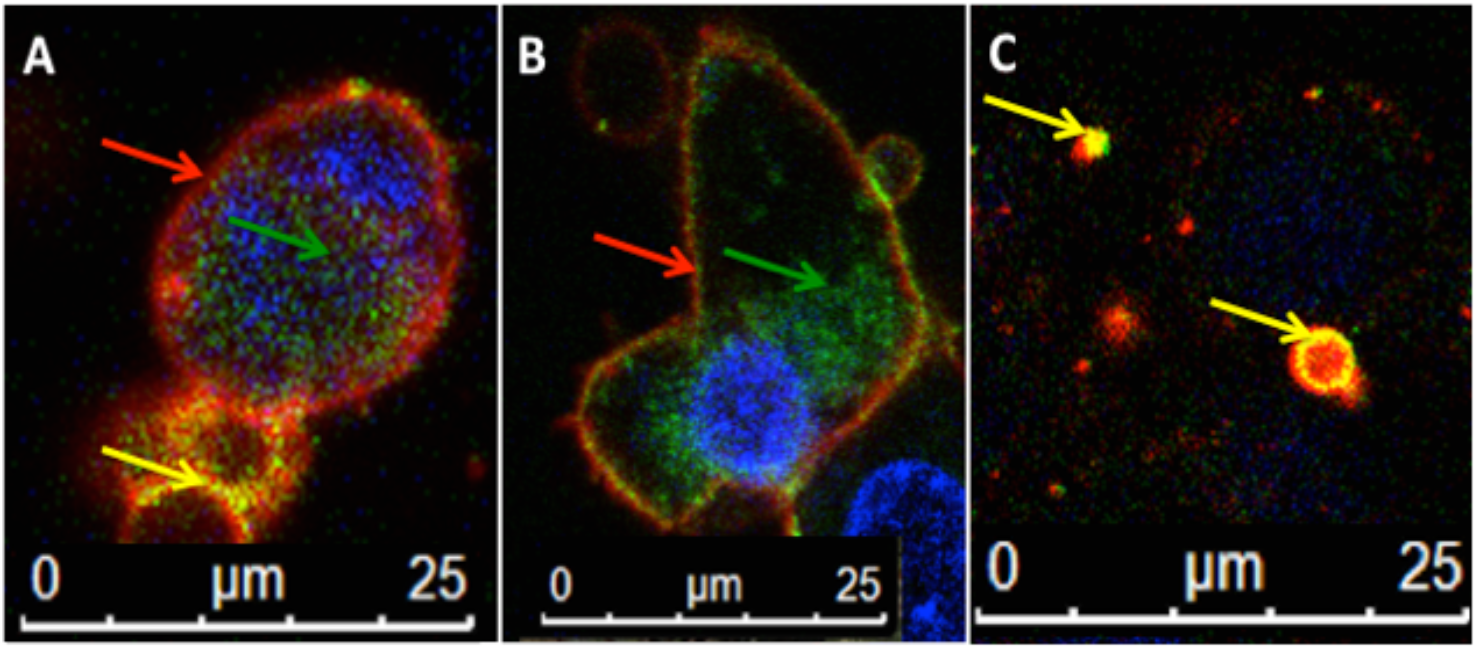
Subcellular localization of GlaS and annexin V. (A and B) Apoptotic 4T1 cells were stained with FITC-GlaS (green arrows) and Cy5-annexin V (red arrows); yellow arrows, co-localization; C) a higher magnification view of possible apoptotic bodies stained as in A and B demonstrating co-staining as an internal control indicating that cellular function seems to be involved in differential staining patterns.

To understand the kinetics of internalization, peroxide-treated cells were co-stained with FITC-GlaS and Cy5-annexin V and imaged using fluorescence microscopy within 10 min. These results demonstrated further distinctions in the staining pattern between the two proteins. For example, Fig 5 shows annexin V strongly staining the surface of one cell while GlaS was internalized in another. Moreover, Hoechst staining revealed that the nuclei of these two cells were also quite different. The cell staining brightly with annexin V had a dense nucleus that might represent a late stage of apoptosis in which chromatin condensation occurs (15). However, GlaS was internalized into cells that displayed a normal diffuse nucleus, suggesting that PS detection occurred in either a potentially healthy cell or a very early apoptotic cell. Moreover, this internalization was rapid, occurring within 10 min. Together with the inability to prevent GlaS binding with excess annexin V (data not shown), these data further support that annexin V and GlaS exhibit different binding and/or uptake properties, and that GlaS may enter the cytosol directly, bypassing receptor mediated endosomal lysosomal uptake.

**Figure 5.**
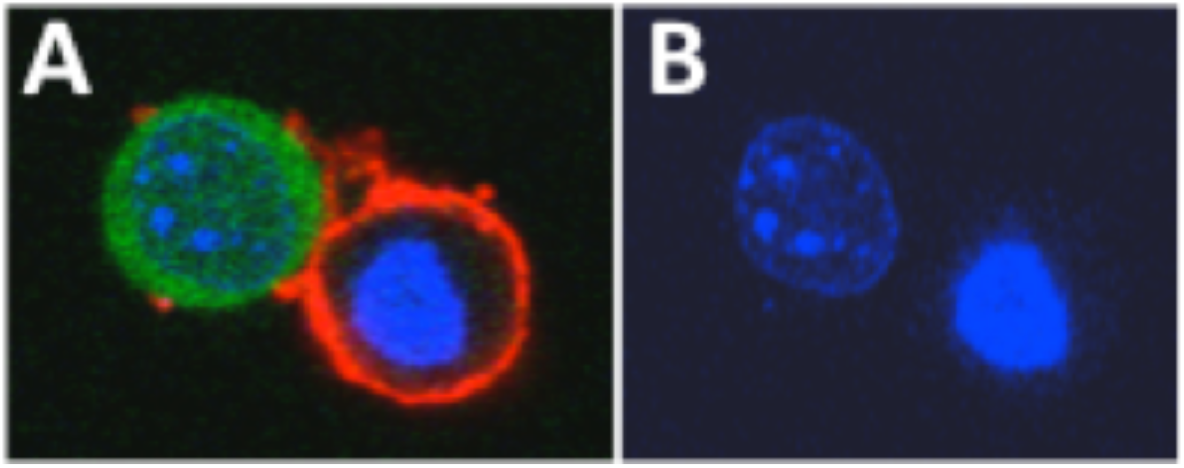
Internalization of GlaS within 10 minutes. Apoptotic 4T1 cells were stained with FITC-GlaS (**green**) and Cy5-annexin V (**red**) and Hoechst (**blue**), and imaged within 10 min of the addition of the proteins. A) Merged image; B) Hoechst nuclear stain alone.

Our observation of co-staining apoptotic bodies (Fig 4C, S1 Fig) prompted a test of GlaS binding to other extracellular vesicles (EV) described to express PS, and a further comparison of this binding to that of annexin V. EVs are expressed from nearly every cell in the body as a means of cell to cell communication and/or waste disposal (16-18) and the study of EVs has become an area of intense research as they have the potential to be used as therapeutic delivery systems (19, 20) and are suspected vehicles for the spread of an array of infectious agents (21, 22). Moreover, stem cells secrete EVs that affect the function of other cells (23) and stem cell-derived EVs are being developed as therapeutics themselves (24). We sought to determine if GlaS could recognize subpopulations of EVs and if this recognition would be distinct from that of annexin V. Because GlaS-stained apoptotic bodies were readily visible among the peroxide-treated cells (Fig. 4C, S1 Fig), we reasoned that EVs could be visualized in the same manner. EVs were isolated from conditioned media of 4T1 cells, followed by staining with GlaS and annexin V. The labeled EVs were imaged by fluorescence microscopy. As seen in Fig 6, the purified EVs do stain with both GlaS and annexin V, similar to the previous studies, demonstrating some overlapping and some distinctive EV staining, respectively. Since EVs were isolated by ultracentrifugation, it is not clear if these differences are due to different subpopulations of EVs (e.g. exosomes, microvesicles, or apoptotic bodies) or if the EVs might be derived from cells in different states (apoptotic or viable, for instance).

**Figure 6.**
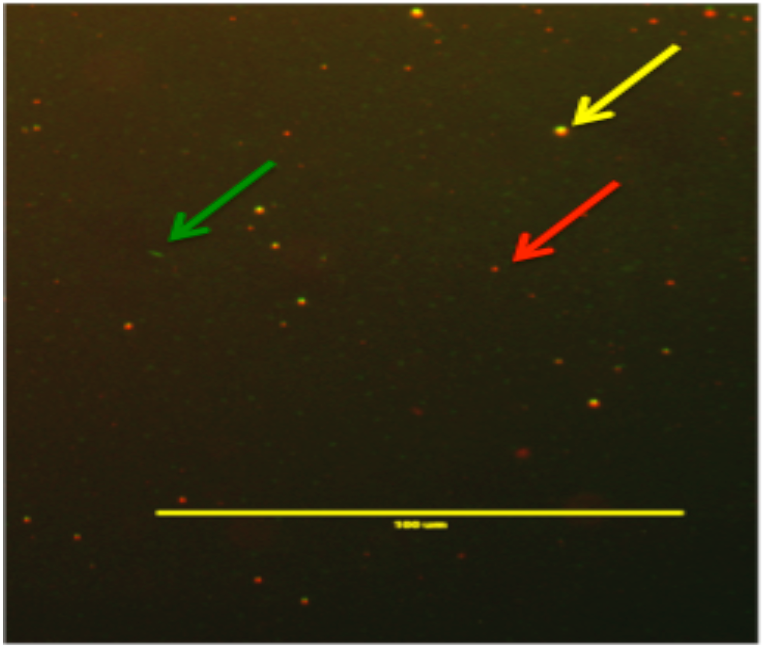
Differential staining of extracellular vesicles by GlaS and annexin V. Extracellular vesicles were prepared by ultracentrifugation from 4T1 cells and stained with FITC-GlaS and Cy5-annexin V. Arrows indicate vesicles staining with annexin V only (red arrow), GlaS only (green arrow) and both proteins (yellow arrow). EVs are smaller than apoptotic bodies with a diameter in the range of 100 nm. Scale bar is 1000 nm.

Multiple different stem cell types stained with both annexin V and GlaS without peroxide treatment (Fig 7) suggest constitutive expression of PS. A subset of murine bone marrow-derived mesenchymal stem cells (MSCs) readily stained with annexin V and GlaS, and the GlaS was internalized into these cells (Fig 7B). Approximately 10% of the MSCs stained in this manner (data not shown). Mouse trophoblast stem cells (TSCs) and murine hematopoietic stem cells (HSCs) exhibited similar staining patterns to MSCs (Fig 7A and C). Interestingly, GlaS uptake into TSCs could occur at 4°C (S2 Fig), suggesting an unusual mechanism of uptake and demonstrating that this process could not be mediated by endocytosis or other energy dependent mechanisms (such as receptor-mediated uptake) that do not occur at a reduced temperature of 4°C. To understand if this recognition of stem cells would be altered if cells were allowed to differentiate, TSCs were grown in the absence of growth factors, a condition which has been shown to cause differentiation into giant cell morphology (25). Interestingly upon differentiation, both GlaS and annexin V could no longer recognize the cells (Fig 8).

**Figure 7.**
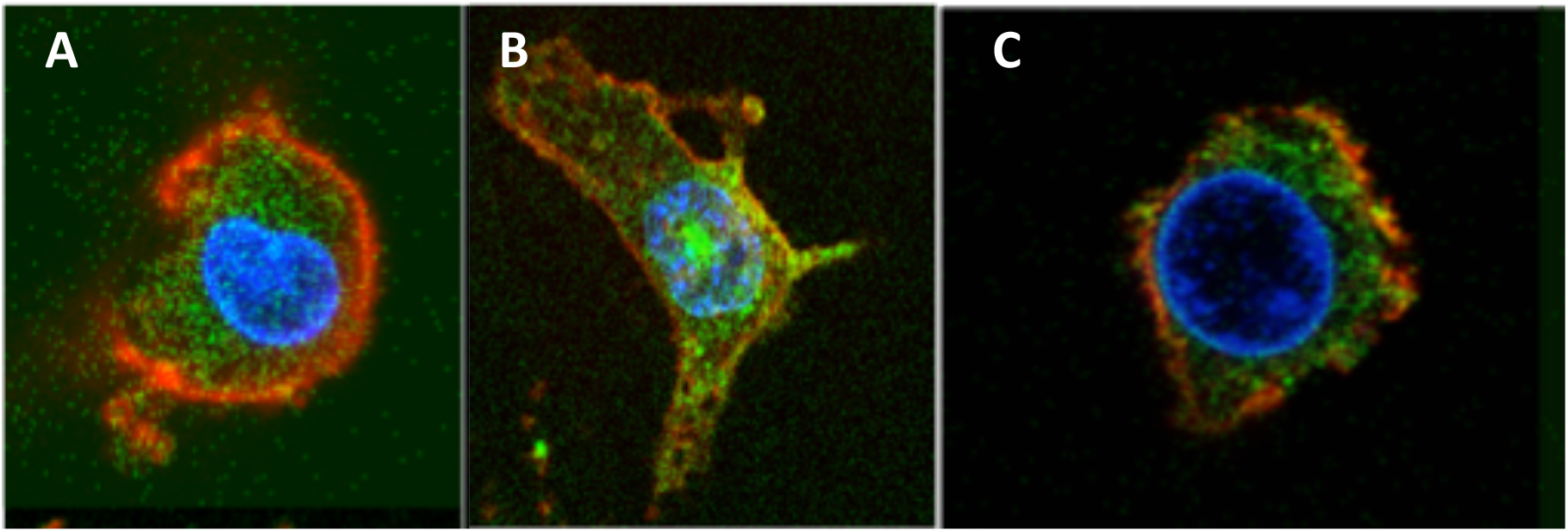
Internalization of GlaS in non-apoptotic stem cells. Live cells were stained with GlaS (green), annexin V (red), and Hoechst (blue) and imaged within 10 min of addition of the stain mixture. A) Trophoblast stem cell; B) Mesenchymal stem cell; C) Hematopoetic stem cell.

**Figure 8.**
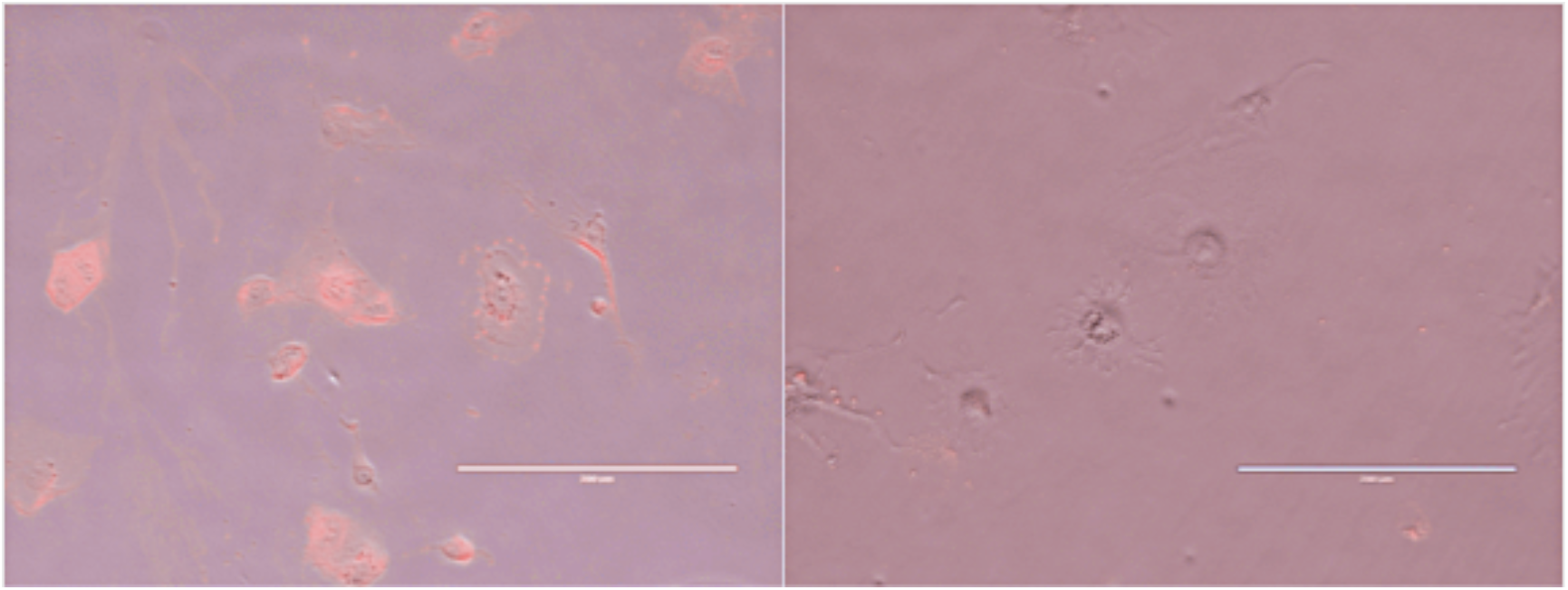
Binding of GlaS to undifferentiated and differentiated TSCs. TSCs grown in the presence (left) and absence (right) of growth factors that prevent differentiation were stained with Cy5-GlaS. Several differentiated giant cells are visible in the center of the right panel.

To further ensure that this process was occurring in stem cells that were not apoptotic, flow cytometry was used to analyze HSCs from mouse bone marrow, a process that is performed the same day as the animal is sacrificed, permitting the identification of live cells by the use of propidium iodide (PI). Supplemental Fig 3 shows the majority of Slam+ HSC stained with GlaS. Dead cells staining positive for PI were excluded from the analysis. Upon examination of the cells by confocal microscopy, GlaS was internalized into live HSCs (Fig S3). Note in this case that annexin stain was detected on the surface, shown in red, but not internalized. An example of a dead HSC is shown in supplemental Fig 3D. Thus, the internalization of GlaS in HSCs occurs in live cells, without the induction of apoptosis.

GlaS is derived from a natural protein, h-protein S, and thus should be non-toxic to cells and nonimmunogenic in humans. To validate the potential of using the GlaS as a platform for PS targeted delivery to cells, we tested the cell viability of TSC over a dose response of GlaS protein (S4 Fig). As predicted, the protein was non-toxic, suggesting that the GlaS protein may be a viable platform for targeting PS expression across a variety of PS expressing cells.

To test for the specific localization of GlaS to sites of PS expression in living subjects, two different models were employed; one to test the ability to detect infection, and the other to detect cancer. For infection, CD1 mice were injected intravenously with bioluminescent *Listeria monocytogenes*. This bacterial pathogen infects many organs including the spleen, in which extensive apoptosis of monocytes and granulocytes occurs (26). At certain times post-infection, spleen is the primary site of bacterial replication and so splenic bioluminescent signals from the bacteria can be correlated with the localization of probes for PS. As seen in Fig 9, GlaS was able to localize to the site of the infection and this signal was absent in uninfected control animals. These results suggest that the GlaS was able to target PS expressing diseased tissue following intravenous administration.

**Figure 9.**
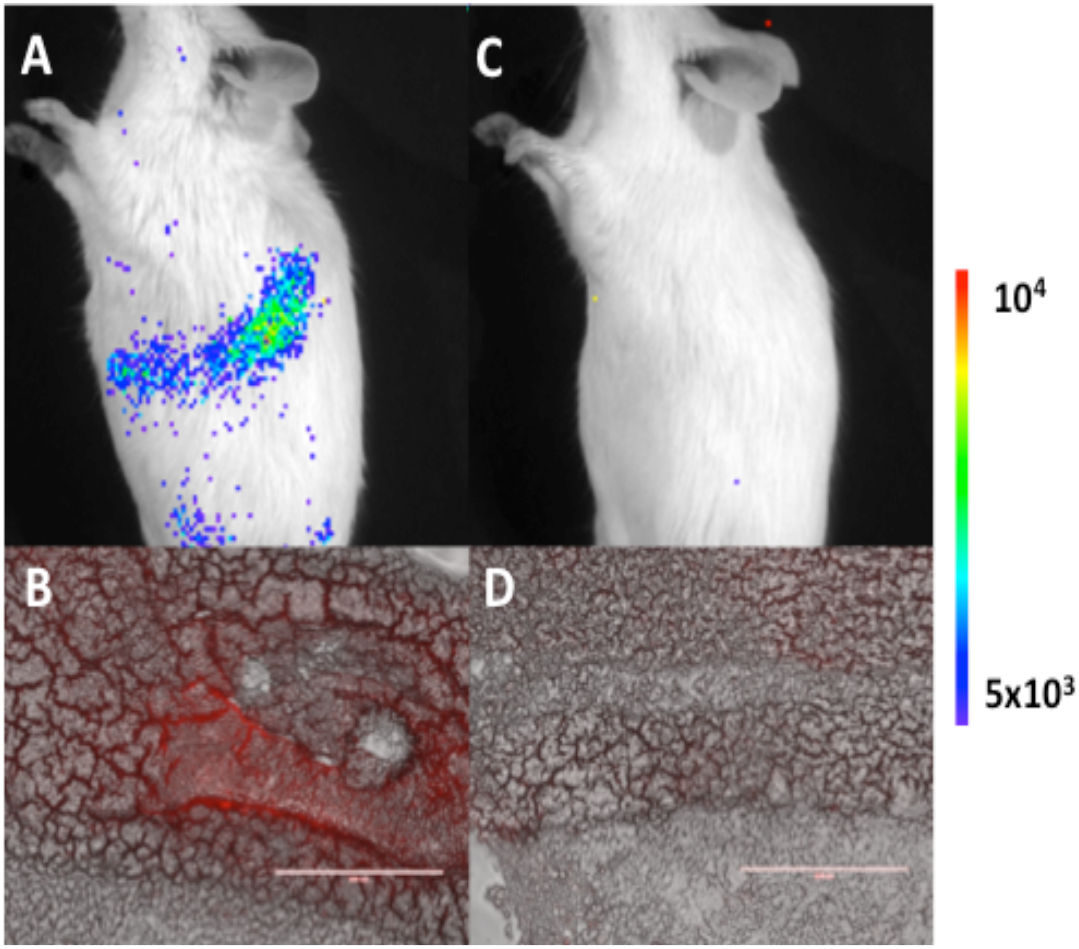
Staining of infected spleen with GlaS. CD1 mice were infected with bioluminescent *Listeria monocytogenes* and imaged on day 2 post infection with an IVIS system. The mice were injected with Cy5-GlaS 30 min before sacrifice, and the spleens removed, frozen in OCT and sectioned for fluorescence imaging. (A) and (C), in vivo bioluminescence images of infected and uninfected mice, respectively. Color bar on right: photons/cm^2^ of the images. (B) and (D), fluorescence microscopy of corresponding splenic sections, merged with phase contrast images.

The localization of fluorescent GlaS to 4T1 tumors with or without doxorubicin treatment was then tested. Doxorubicin has been shown to enhance PS externalization (27), and following Cy5-GlaS administration, GlaS localization was identified (Fig 10). Areas of intense Cy5-GlaS staining were observed in tumors of doxorubicin-treated animals, whereas more modest signal was observed from the untreated tumor sections. These results suggest that GlaS was able to target PS expressing tumors following intravenous injection and that this signal was intensified by the treatment of doxorubicin.

**Figure 10.**
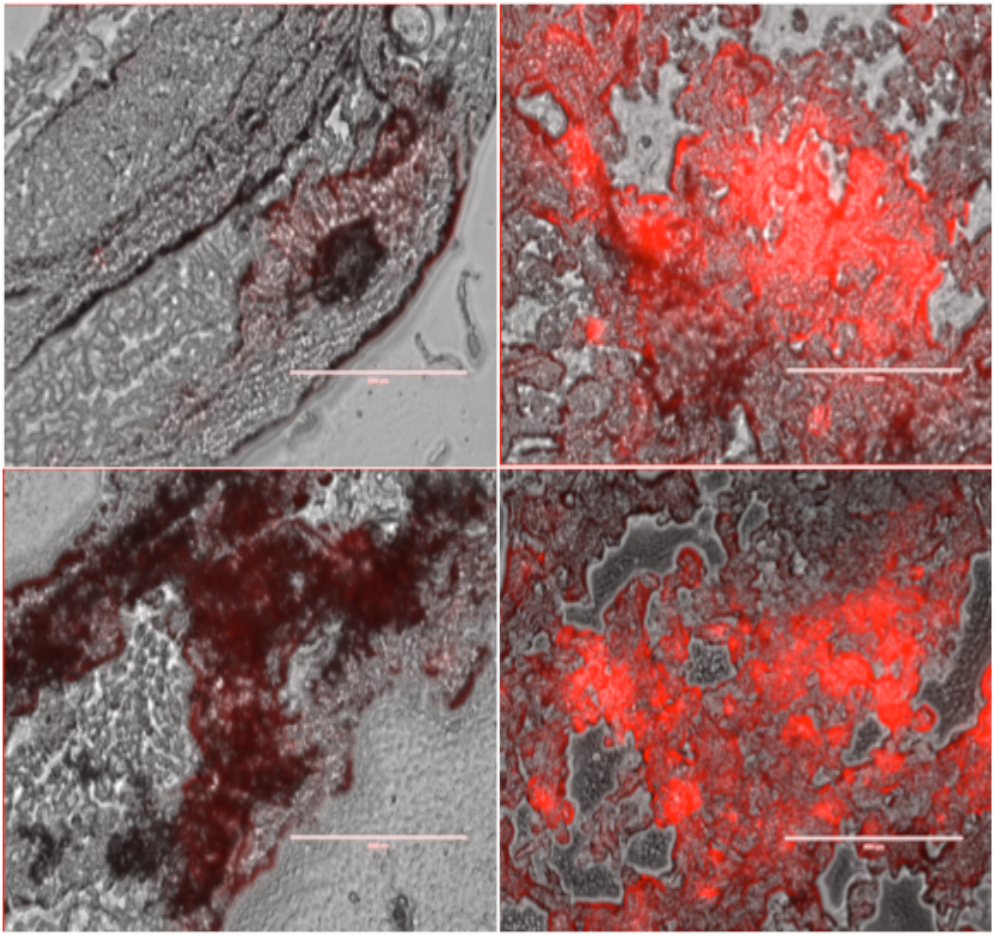
Localization of Cy5 GlaS to tumors treated with doxorubicin. Mice implanted with 4T1 breast cancer tumors were treated with doxorubicin (right panels) or left untreated (left panels). 24 hours post treatment the mice were injected intravenously with Cy5-GlaS and sacrificed 30 min later. The tumors were removed, frozen, and sectioned for fluorescence microscopy. Merged Cy5/phase contrast images from four different mice are shown.

## Discussion

Under physiological conditions, externalized PS functions as a dominant and evolutionarily conserved immunosuppressive signal that promotes tolerance and prevents local and systemic immune activation (2). For example, PS plays a vital role in tissue homeostasis by aiding in the removal of dying cells while suppressing potentially detrimental inflammatory responses in the process. Pathologically, the innate immunosuppressive effect of externalized PS has been hijacked by numerous infectious agents (viruses, microorganisms, and parasites) to facilitate infection and establish infection latency (22, 28). PS exposure is also dysregulated in diseases, such as cancer, where exposure on the tumor cell and in the tumor microenvironment antagonizes the development of tumor immunity (3). Importantly, and perhaps not surprisingly, this immunosuppressive effect can be extended from the diseased cell and organism to other healthy cells or organisms through the release of PS-coated EVs (22, 29). Consequently, agents targeting PS could have significant value as cancer and infectious disease therapeutics and/or diagnostics.

The idea of targeting PS as a therapeutic and diagnostic is not a new concept, especially in cancer. However, to date it has not been possible to harness the full potential of PS for delivery of therapeutics and also as a diagnostic. Annexin V has been tested in various clinical trials as a diagnostic agent (30) and attempts have been made to translate annexin V into a therapeutic entity but have been hampered by 1) the need for annexin to trimerize to interact with PS (31) and 2) it’s cell surface localization, limiting potential to deliver therapeutics to the outside of the cell. The antibody, bavituximab, a chimeric monoclonal antibody that targets the PS-binding serum protein β2 glycoprotein-1 (β2GP1) (32), failed a Phase III clinical trial in non-small cell lung cancer patients in combination with docetaxel (33, 34). One potential limitation of bavituximab was its inability to internalize and additional work has been initiated to graft naturally occurring PS-targeting agents to Fc fragments to take an approach mimicking an antibody drug conjugate (35). However, this approach still faces the challenges of current antibody drug conjugates being efficient recognition, internalization and release of the drug while traversing the endosomal and lysosomal pathway (36). In general, new tools are needed to harness the potential of PS targeting for the treatment of diseases, including cancer.

In this manuscript we describe the characterization of the GLA domain, a motif shared between proteins involved in blood homeostasis (Fig 1), and our efforts to exploit its natural PS targeting as a tool for imaging, diagnosis, or therapeutic treatment of diseased cells and their associated EVs. As a naturally occurring protein motif found in the body, the delivery platform should have reduced immunogenicity potential and the advantage of being non-toxic (Fig S4). We were able to confirm that the GLA domain, as isolated, was able to recognize cell surface exposure of PS (Figs 2, 3, 7) and that this recognition was selective for cell surface PS expression (Figs 2 and 8). Interestingly, the interaction between the PS-binding proteins annexin V and GlaS with PS displayed some significant differences. For example, in PS-expressing cells (Figs 3, 4, 5, 7) and EVs (Fig 6) stained with fluorophore-tagged GlaS and annexin V revealed that there were cells and EVs that shared binding of both annexin V and GlaS and also cells and EVs that were only recognized by either annexin V or GlaS. Vesicles of a size consistent with apoptotic bodies stained with both proteins, suggesting that cellular functions may determine differential binding (Fig 4). One hypothesis is that the annexin V sees only cells that are fully committed to apoptosis as evidenced by the nuclear changes, whereas GlaS may detect cells that have started the apoptotic journey but are not fully committed to the same. Importantly, and in contrast to annexin V, GlaS was able to internalize into PS-expressing cells (Figs 3, 7). This ability to recognize, rapidly internalize (less than 10 min) in an energy-independent fashion into the cytoplasm of a variety of PS-expressing cells (*ex vivo* stem cells, infected cells, cancer cells) and interact with PS-expressing EVs has potentially important therapeutic and diagnostic implications across an array of diseases and biological functions.

While annexin V and GLA domain both bind PS, there appear to be differences in their interaction with PS exposed on the surfaces of cells and particles. These differences are significant and suggest advantages for the GLA domain over annexin V as a basis for clinical applications. First, the GLA domain appears to detect PS earlier in the apoptotic process than annexin V. Secondly, upon cell surface PS binding, Gla is internalized, whereas annexin V remains on the cell surface. This internalization is unusual in that it is rapid (within 5 min exposure to the cell) and can occur at 4°C suggesting an ATP-independent non-endocytic pathway for direct entry into the cytosol. This internalization is independent of cell type and is also observed in cultured and primary stem cells, with the displayed PS on their cell surface that was susceptible to detection and internalization of the GLA domain. This was not observed with annexin V across an array of stem cell types. Lastly, annexin V and the GLA domain demonstrate differential interaction with extracellular vesicles (EVs). These results indicate that PS-binding by GLA domains is distinct from annexin V in ways that suggest that engineered recombinant GLA domains may serve as unique PS-targeted intracellular delivery vehicles for therapeutic, imaging, diagnostic, and cell engineering purposes by targeting cells in specific physiologic state marked by PS expression.

Stem cells hold great potential for the treatment of an array of diseases but generally, require gene manipulating tools (e.g. CRISPR/Cas9, RNAi, ZFN, or TALEN) to enable their full therapeutic potential. Novel delivery methods are needed that eliminate the toxic effects or safety problems associated with the current methods (electroporation, transfection, virus-based methods). The GlaS platform may address many of these liabilities given that it is non-toxic to stem cells (Fig S4), and as a protein, possesses a transient lifetime in the cell. Tools that directly deliver attached payloads to the cytoplasm of cancer stem cells for elimination, or to normal stem cells for directed reprogramming, would have tremendous value as the foundation for targeted therapies for a wide variety of diseases.

As a tool for differential staining of EVs, GlaS may help in the understanding of their biology and in aiding diagnosis of a variety of diseases. EVs contain much of the information that is available in liquid biopsies, and the ability to capture and/or identify EVs from different cell sources has potential in diagnosis, prognosis, and therapeutic guidance (37). These nanosized particles transmit signals determined by their protein, lipid, nucleic acid and sugar content, and the unique molecular pattern of EVs dictates the type of signal to be transmitted to recipient cells. The composition of EVs is not random and each EV-cargo delivers specific molecular messages (38) which makes them attractive as potential vaccine sources. However, their small sizes and the limited quantities that can usually be obtained from patient-derived samples pose a number of challenges to their isolation, characterization, and engineering. In addition, EVs are ‘pre-programmed’ with selected cargoes and cell-specific targeting moieties, which may not necessarily overlap with their intended therapeutic application. Given the reported expression of PS on EVs and the confirmation that this is seen by GlaS and differentially from annexin V, GlaS may have a potential roll here as a means for differential isolation and manipulation of EVs and is an area of active current research.

Lastly, it is important to note that infectious pathogens have evolved many elegant strategies to manipulate host cells to further their pathogenic potential, and this includes adopting the cloaking mechanism of apoptotic mimicry, defined by the exposure of PS on the pathogen surface (28). Thus, therapeutic targeting of PS may represent an opportunity to try to realize new broad-spectrum anti-infectious agents and has important ramifications as a potential frontline therapy for intervention in outbreaks where the infectious disease entity is not initially known. Our experiments demonstrating the *in vivo* PS detection capability of this platform for both cancer and bacterially-infected cells (Figs 9, 10) suggest that it has potential for clinical translation for an array of disease states which display cell surface PS as a hallmark of disease.

## Conclusions

In summary, we have identified a novel platform for PS-mediated targeted delivery of therapeutics to an array of diseases (*i*.*e*., infectious diseases, cancer) and cellular targets (*i*.*e*., stem cells, extracellular vesicles). Targeting a host cell change fundamental to disease instead of a disease specific protein has significant ramifications. For example, envelope viruses (*e*.*g*. Influenza, COVID-19) (28) have been described as having PS on the envelope of the infectious virus and on the surface of the infected cell. Targeting via a virally induced change fundamental to the disease biology would be a new paradigm that avoids pitfalls to current viral treatments, where viral mutation and protein plasticity creates viral escape mutants that thwart antiviral and vaccine efforts. And this extends to oncology, developing a pan anti-cancer approach for the treatment of an array of cancers. Consequently, future studies will focus on defining and optimizing the payload capacity and its application to areas of unmet medical need.

## Acknowledgements

We wish to thank John Teare and Doug Schneider for their thoughtful review of the manuscript. This work was funded in part by the generous support of the James and Kathleen Cornelius endowment to MSU (CHC).

