## Supplemental data for "Gla-domain mediated targeting of externalized phosphatidylserine for intracellular delivery"

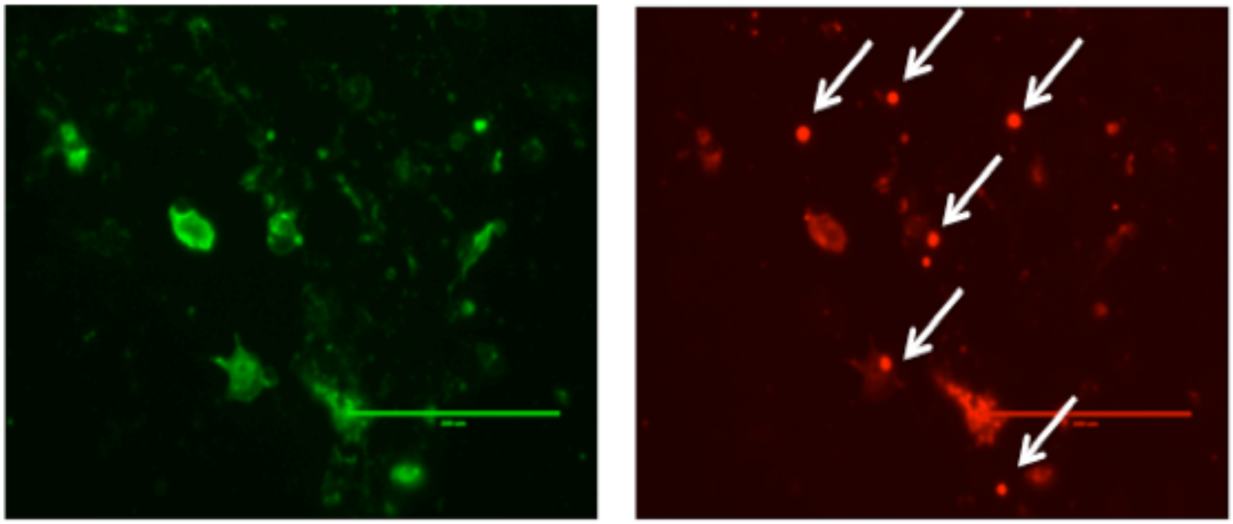

**Supplementary Figure 1. Staining of apoptotic COS-1 cells with GlaS and annexin.** Cells were treated with t-BHP and stained with FITC annexin (**green** - left) and Cy5 GlaS (**red** - right). Arrows indicate subcellular structures presumed to be extracellular vesicles.

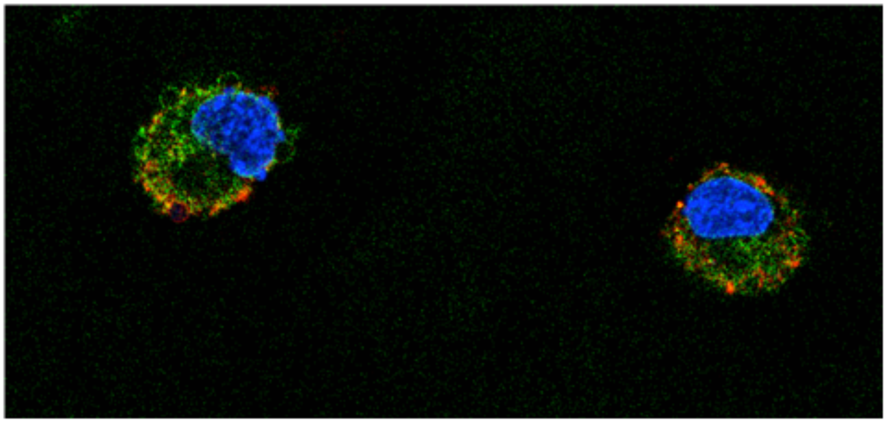

**Supplementary Figure 2. Entry of GlaS into the cytosol at 4C.** TSC cells at 4C were stained with FITC GlaS (**green** - left) and Cy5 annexin (**red** - right) and imaged 5 minutes later while still cold.

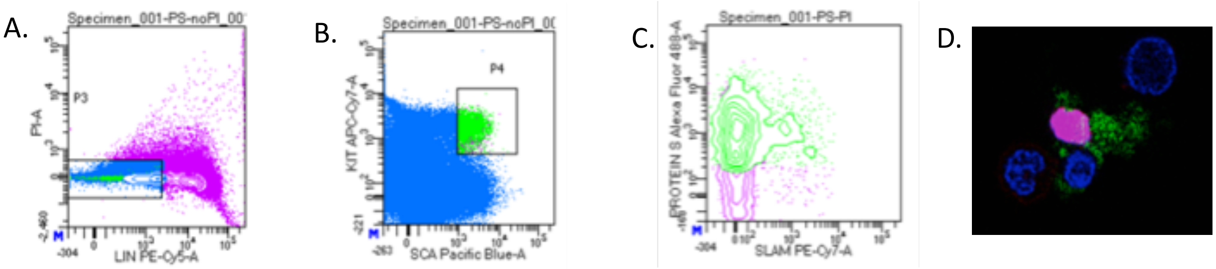


**Supplementary Figure 3. Flow cytometric analysis of HSCs, and PI staining pattern of dead HSCs.** Lineage-negative, SCA-1/c-kit staining cells from mouse bone marrow: **A**) Absence of staining for hematopoietic lineages and **B**) staining of c-kit and SCA1 defines the population of HSC, shown in green. **C**) GlaS staining of long-term HSC. HSC were isolated and stained with FITC GlaS. SLAM pattern was determined with Cy7 (x-axis). **D**) An example of a dead cell (exhibiting PI staining of the nucleus in pink) among three live cells (blue nuclei); also compare to live cell in Figure 7c.

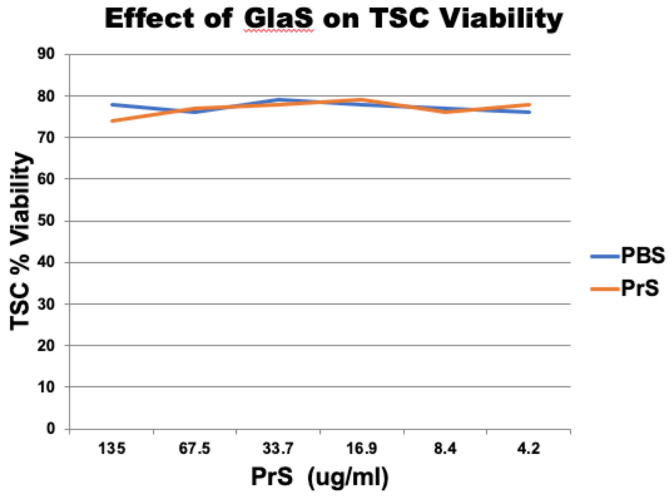


**Supplemental Figure 4. Toxicity of GlaS in cultured TSCs**. TSCs were treated with GlaS at the indicated concentrations and viability was measured by trypan exclusion after 30 minutes. Percent was determined using a Cellometer.
